# Microglia morphology in the developing primate amygdala and effects of early life stress

**DOI:** 10.1101/2024.08.15.608133

**Authors:** Dennisha P. King, Miral Abdalaziz, Ania K. Majewska, Judy L. Cameron, Julie L. Fudge

## Abstract

A unique pool of immature glutamatergic neurons in the primate amygdala, known as the paralaminar nucleus (PL), are maturing between infancy and adolescence. The PL is a potential substrate for the steep growth curve of amygdala volume during this developmental period. A microglial component is also embedded among the PL neurons, and likely supports local neuronal maturation and emerging synaptogenesis. Microglia may alter neuronal growth following environmental perturbations such as stress. Using multiple measures, we first found that microglia in the infant primate PL had relatively large somas, and a small arbor size. In contrast, microglia in the adolescent PL had a smaller soma, and a larger dendritic arbor. We then examined microglial morphology in the PL after a novel maternal separation protocol, to examine the effects of early life stress. After maternal separation, the microglia had increased soma size, arbor size and complexity. Surprisingly, strong effects were seen not only in the infant PL, but also in the adolescent PL from subjects who had experienced the separation many years earlier. We conclude that under maternal-rearing conditions, PL microglia morphology tracks PL neuronal growth, progressing to a more ‘mature’ phenotype by adolescence. Maternal separation has long-lasting effects on microglia, altering their normal developmental trajectory, and resulting in a ‘hyper-ramified’ phenotype that persists for years. We speculate that these changes have consequences for neuronal development in young primates.

**Significance Statement:** The paralaminar (PL) nucleus of the amygdala is an important source of plasticity, due to its unique repository of immature glutamatergic neurons. PL immature neurons mature between birth and adolescence. This process is likely supported by synaptogenesis, which requires microglia. Between infancy and adolescence in macaques, PL microglia became more dense, and shifted to a ‘ramified’ phenotype, consistent with increased synaptic pruning functions. Early life stress in the form of maternal separation, however, blunted this normal trajectory, leading to persistent ‘parainflammatory’ microglial morphologies. We speculate that early life stress may alter PL neuronal maturation and synapse formation through microglia.

---

During infancy, human and nonhuman primates solidify a permanent foundation for their cognitive, social, and emotional health. One structure critical for social and emotional development is the amygdala (Bachevalier et al., 1999; Sanchez et al., 2001; Amaral et al., 2003; Gee et al., 2013; Tottenham, 2014), which detects and processes salient sensory cues in awake human and nonhuman primates (Wang et al., 2014; Wang et al., 2017; Grabenhorst et al., 2019; Morrow et al., 2020). Lesions and environmental factors can impact the maturation of the amygdala resulting in life-long consequences for affective behavior (Schumann et al., 2011; Payne and Bachevalier, 2019).

The amygdala volumetrically expands between infancy and adolescence in both monkeys and humans (Giedd et al., 1996; Uematsu et al., 2012; Hu et al., 2013; Goddings et al., 2014; Schumann et al., 2019). Despite neuroimaging evidence for volumetric growth of the amygdala, few studies have examined the cellular basis for this expansion. Prior studies in primates as well as in rodents, indicate that the amygdala’s mature neuron numbers increase between birth and adolescence. In humans, dendritic complexity also increases (Rubinow and Juraska, 2009; Weir et al., 2017; Avino et al., 2018; Sorrells et al., 2019; McHale-Matthews et al., 2023).

A potential source of increasing numbers of mature neurons in the amygdala during early life is the paralaminar nucleus (PL). The PL is a repository of immature glutamatergic neurons that persist postnatally. Between infancy and adolescence, the ratio of mature to immature neurons in the PL increases, suggesting active maturation (Avino et al., 2018; Sorrells et al., 2019; Chareyron et al., 2021; McHale-Matthews et al., 2023). In humans, other amygdala nuclei also gain mature neurons in the first decades of life (Avino et al., 2018), leading to the idea that neuronal maturation, and possibly migration, from the PL, is a potential mechanism for cellular expansion not only in the PL, but also other amygdala nuclei.

One major question is: what neural processes support PL cellular maturation between infancy and adolescence? Emergent excitatory activity from afferent terminals is required for post-synaptic neuron development, stimulating the growth of dendritic spines, elaboration of dendritic arbors, and synapse formation (Engert and Bonhoeffer, 1999; Harris, 1999; Evers et al., 2006; Elston and Fujita, 2014; Oga et al., 2017). Thus, excitatory inputs that arrive in the amygdala postnatally may also shape PL neuron maturation (Cressman et al., 2010; Arruda-Carvalho et al., 2017; Murray and Fellows, 2021). Environmental influences such as sensory changes and stress can therefore influence spinogenesis and synaptogenesis in the developing brain, presumably through effects on this process (Vyas et al., 2002; Mitra et al., 2005; Roozendaal et al., 2009; Block et al., 2022).

Microglia are immune cells but also aid in neural network assembly in the developing brain through synaptic remodeling (Paolicelli et al., 2011; Hammond et al., 2018; Whitelaw et al., 2023). Microglial phagocytosis of weak synapses is essential during circuit formation (synaptogenesis) because it maintains and strengthens the remaining synapses, thereby enhancing neural transmission (Schafer et al., 2012; Zhan et al., 2014; Whitelaw, 2018). For example, classic studies of light deprivation in juvenile mice show increased microglial engulfment of weakened synaptic elements including synaptic clefts, resulting in repatterning of the visual field (Tremblay et al., 2010; Tremblay and Majewska, 2019). Despite its high degree of neuronal plasticity, microglia in the PL have not been previously characterized in a macaque.

In view of the extensive literature confirming that early environmental influences represent a major risk factor for the subsequent development of several psychopathologies (Kessler et al., 2003; Cameron et al., 2017; Catale et al., 2020), and the important role of microglia in establishing circuitry (Wei et al., 2015; Delpech et al., 2016), we asked whether a potent developmental event would alter microglial morphology, a potential sign of altered synaptogenesis. Using classic morphologic criteria, we investigated both normal PL microglial development, as well as the effects of modified maternal separation in infancy and adolescence.

## Methods

### Animals

The twenty-three primates (*Macaca mulatta)* used in this study were born and reared at the University of Pittsburgh (N= 19 female, N=4 male). All animals were housed in group-rearing pen environments composed of 4-5 primates ranging in age from adolescence to adulthood. Animals were divided into cohorts, exposed to maternal separation, and behaviorally assessed. All experiments were approved by the University of Pittsburgh Animal Care and Use committee and were in accordance with guidelines issued by the National Institute of Health. Following sacrifice, brains were transferred to the University of Rochester for assessments. Tissue from some of these animals was previously used for amygdala gene expression and neuronal cell counting (Sabatini et al., 2007; de Campo et al., 2017; McHale-Matthews et al., 2023).

### Maternal Separation Paradigm

The maternal separation protocol has been extensively described in previous publications and is described here in brief. For full details of the animal husbandry and the modified maternal separation paradigm, see (Sabatini et al., 2007; deCampo and Fudge, 2012; Cameron et al., 2017; McHale-Matthews et al., 2023). Two age cohorts, an infant (3 months at sacrifice) and an adolescent (4-5 years at sacrifice) were exposed to similar protocols beginning at birth **(Fig. 1)**. In the cohort sacrificed at 3 months, all animals were female (n=12). In the cohort sacrificed in adolescence, there were 3 males, and 8 females (n=11). All infants shared a single cage with their mother during the first week of life (per typical animal husbandry protocols) and were then randomly assigned to one of the following experimental groups: “1-week separated (1-WS),” “1-month separated (1-MS),” or “maternally reared (MR)”. The randomization to 2 separation groups was based on extensive research on the impact of timing of environmental exposures on behavioral outcomes in monkeys (Sabatini et al., 2007; Cameron et al., 2017).

**Figure 1.**
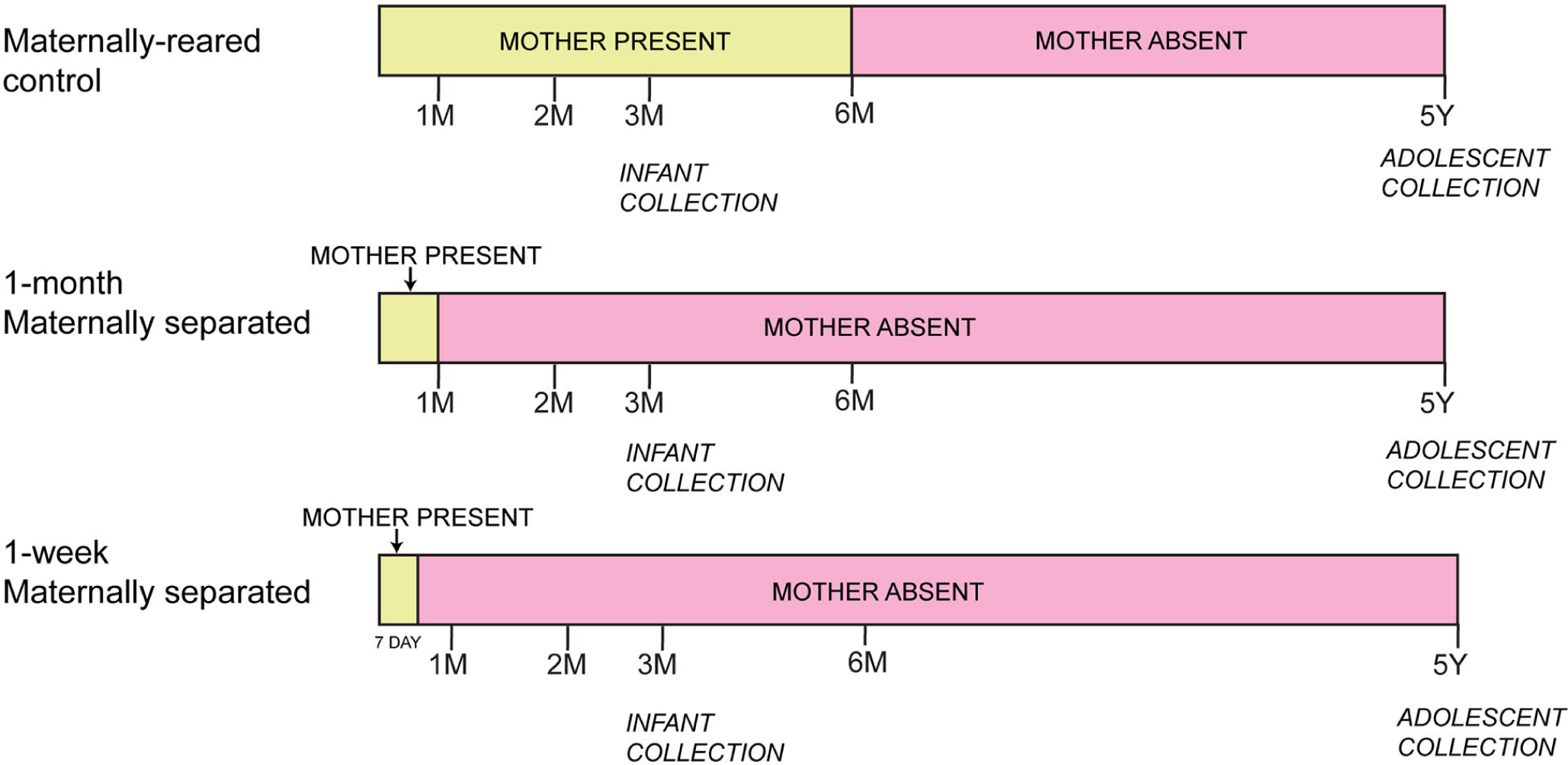
General design of the maternal separation paradigm in infants and adolescent animals. See Methods for details.

At the beginning of the study, only 1 experimental infant and its mother were present in each group-rearing pen, with 3 nonexperimental monkeys of varying ages (juvenile to adolescent in age). When the mother was present, her status was therefore the most dominant member of the group, to insure there was no added stress on the mother. Maternally reared (MR) and 1 month separated (1-MS) animals were introduced to a group-rearing pen environment with their mother present from the beginning of week 2. The 1-WS group had their mother removed but was left in the single cage and taught to bottle feed for a 5–7-day period (*ad libitum* Similac with Iron baby formula; Abbott Laboratories, Columbus OH). They were then placed in the pen at week 2 of age and a hutch with a door small enough for the separated infant to transgress, but too small for other animals to enter, was placed in the pen and bottles were made available to the infant in this cage. A soft, cotton-stuffed toy was placed in the cage to provide contact comfort. In contrast. 1-MS animals were integrated into a group-rearing pen environment with their mother present for weeks 2-4. At the end of week 4, the 1-MS animals were separated from their mothers and trained to bottle feed in a single cage adjacent to the pen for a one-week period and then re-introduced to the same group-rearing environment for weeks 5-12. We analyzed the two separation groups individually to investigate possible timing effects of maternal separation. For the infant cohort, the MR group had their mother present in the pen for the 12 weeks of the study. For the adolescent cohort, maternally reared (MR) animals continued with the mother in the group-rearing pen through 6 months of age, since this is the typical point at which macaque mothers leave their infant and form a consort with a male macaque for the next mating season (Cameron et al., 2017).

For the adolescent cohort, monkeys were reared until they were 4.43±0.16 years of age in the stable group pens which they were placed in at 2 weeks of age. They were then regrouped into new social groups, consisting of one 1-WS, one 1-MS, and one 6-MS to study their behavioral response to entering a new social group. Adolescent monkeys were sacrificed 8.95±0.31 months after regrouping, when they were 5.14±0.17 years of age.

### Tissue Collection and Processing

Brain tissue from the infant cohort was harvested at week 12. Brain tissue from adolescent animals was harvested at 4-5 years. All animals were first anesthetized with ketamine HCl (10 mg/kg, i.m. injection), then deeply anesthetized with sodium pentobarbital (30 mg/kg, i.v.). A transcardial perfusion surgery protocol was followed, using: 1 liter of ice cold 0.9% NaCl solution containing 2% sodium nitrite and 5000 IU heparin. The brain was removed, hemisected on the mid-sagittal plane. The left hemisphere used in this study was cut in coronal blocks and immersion fixed (Sabatini et al., 2007; de Campo et al., 2017). Blocks were sectioned coronally on a freezing sliding microtome at 40μm thickness and the rostral to caudal extent of the amygdala was saved in serial compartments.

### Histology for fixed tissue

#### Immunocytochemistry

1:12 sections from adjacent compartments were chosen to ensure rostral to caudal sampling of the entire amygdala, yielding approximately 7-9 sections per case. Protocols for doublecortin (DCX) and ionized calcium binding adaptor 1 (IBA1) immunolabeling were first established in control animals that were not part of this study (Fudge et al., 2012). DCX protein identifies immature neurons of the PL, distinguishing them from overlying basal nucleus (de Campo et al., 2017; Page et al., 2022; McHale-Matthews et al., 2023). IBA1 is a cytoplasmic protein found in macrophage-derived cells such as microglia (Ito et al., 1998). Cases were immunolabeled in batches in a blinded counter-balanced fashion as follows.

Tissue was rinsed in PB with 0.3% Triton-X (PB-TX) overnight. The next day, the sections were first treated with an endogenous peroxidase inhibitor for 5 minutes, and then rinsed in PB-TX for a total of six rounds, 15 minutes each round with the replacement of fresh PB-TX. After the rinses, sections were pre-incubated for 30 minutes in 10% normal donkey serum blocking solution with PB-TX (NGS-PB-TX). Sections were then incubated in primary antisera to DCX (1:15000, Abcam, rabbit) or IBA1 (1:5000, FujiFilm, rabbit) at 4°C on a rocker for four nights. Sections were again rinsed, blocked with 10% NGS-PB-TX, and incubated for 40 minutes in anti-rabbit biotinylated secondary antibody. After secondary antibody incubation, and sequential rinses over 1.5 hours, sections were incubated in an avidin-biotin complex for one hour (Vectastain ABC Elite; Vector Laboratories), then visualized with 3,3’-Diaminobenzidine (DAB).

All immunolabelled sections were rinsed overnight to reduce background staining, and then mounted onto gelatin treated slides from 0.1M PB solution and left to air dry for 2-4 weeks. Once dry, DCX-labeled sections were additionally dehydrated, rehydrated, and then counterstained lightly with cresyl violet (Chroma-Gesellschaft; West Germany). All sections were cover slipped with DPX Mounting Medium (Electron Microscopy Sciences; Hatfield, PA).

### Analysis

#### Location of regions of interest (ROIs)

All slides from each case were coded to ensure that investigators were blind to age and maternal separation condition. For each case, we assessed 4-5 evenly spaced slides matched for rostral to caudal level across animals. The PL was identified in DCX/Nissl labeled sections using brightfield microscopy with a 2x objective **(Fig. 2A).** Intensely labeled DCX-positive immature neurons are seen in the laminar PL structure ventral to the basal nucleus, and a contour was drawn around this region using mapping software (Olympus AX70 microscope interfaced with Stereoinvestigator via a video CCD, Microbrightfield, Williston, VT). Adjacent IBA1-labeled sections were captured at the same magnification, and contours were aligned onto matching landmarks such as traced such as blood vessels and fiber tracts to localize the PL (**Fig. 2B**). Within the overlaid contour, the IBA1-labeled cells in PL in three ROIs were photographed using a 40x objective. Images were initially collected in medial, central, and lateral ROIs for each section. However, in preliminary analyses (not shown), we determined that there were no statistical differences in microglial density, spacing, or soma size across these regions in any animal. Therefore, data from these locations was pooled for PL microglia density counts, soma size, and spacing index values.

**Figure 2.**
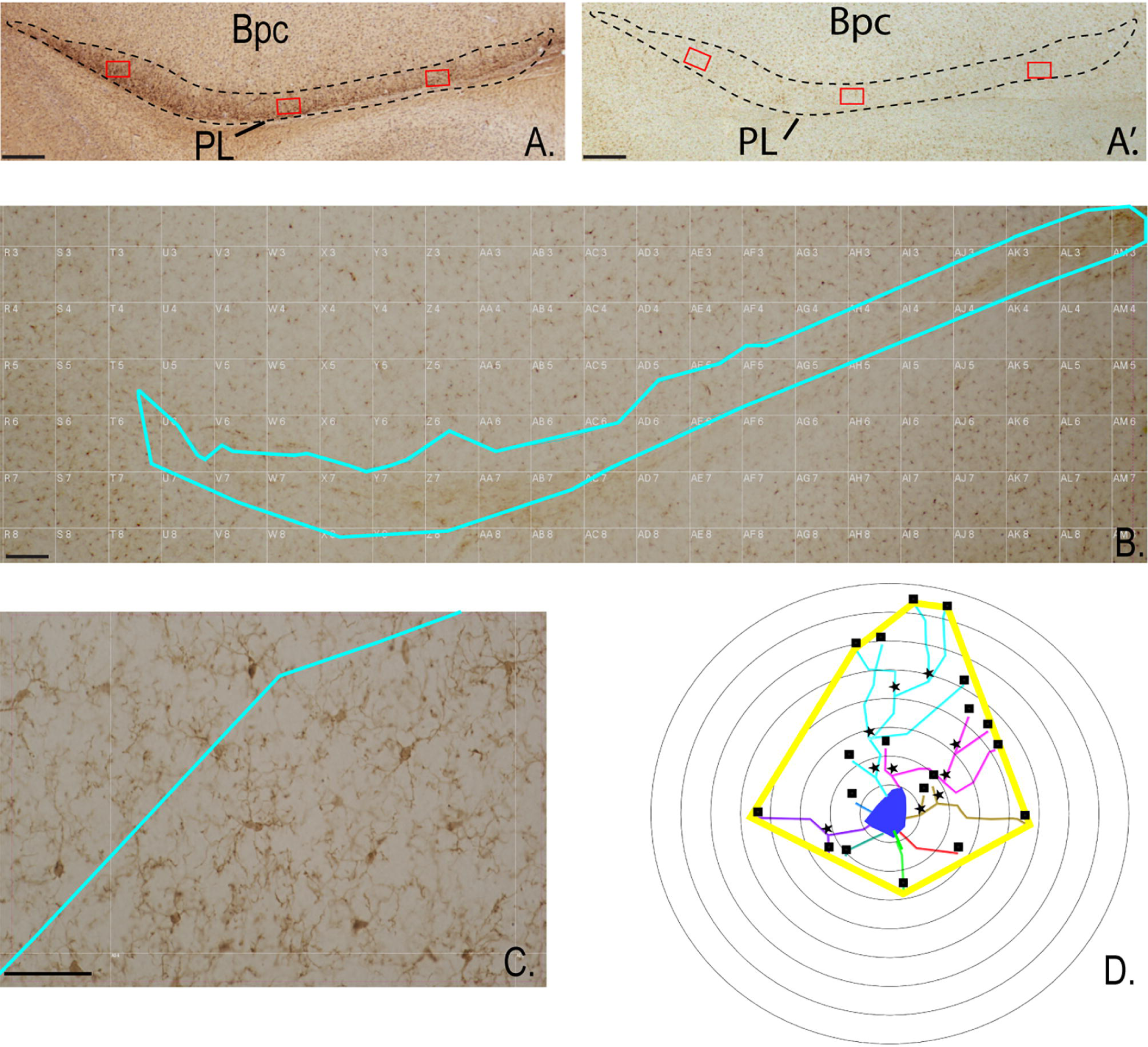
A. Low power image of dense DCX-IR which defines the PL, also shown with contour3. A’. Adjacent section immunoreacted for IBA1, with the same PL contour superimposed. White ROI selection gird boxes for collection of density, spacing index, and soma size are shown. Bars =25µm B. 2x view of grid used for localizing central PL ROI for microglia morphology sampling. Bar= 25µm C. 40x image of the region in B. Bar=25µm D. Schematic of representative Sholl measurements. Blue delineates the cell soma, individual processes originating at the cell soma are shown in different colors, while stars mark nodes and squares mark endpoints. The thick yellow line shows the convex hull.

#### Microglial Morphology Data Collection

##### Microglial density

During normal development, microglia density increases in many brain regions between infancy and adolescence (Menassa et al., 2022) and can also be altered by early life stress (Reemst et al., 2022). Therefore, we calculated the area of the PL by using the freehand tool in Fiji/Image J to re-trace the previously outlined contour of the PL. All microglia within each ROI were selected using the multi-point tool, which also automatically records the X and Y position of each selected microglia. Microglia density was calculated as the average total number of microglia per ROI area.

##### Spacing index

The spacing of the microglia was calculated to determine the nearest neighbor distance between individual microglia, corrected for density. During normal development microglia are typically evenly distributed, and in general, spacing does not significantly differ depending on age (Savage et al., 2019). The ‘spacing index’ (or ‘clustering index’) is calculated as: (average nearest neighbor distance)^2^*microglial density. The spatial coordinates captured (above) were entered into a customized Matlab code (Mathworks) where the distance between each microglia and its nearest neighbor was determined and averaged for all microglia per ROI.

##### Soma area

The normal developmental transition from infancy to adolescence is often marked by decreased microglial soma size (Savage et al., 2019). However, in pathological conditions this transition is broadly impacted (Reemst et al., 2022). Therefore, we examined a total of 80-100 microglia per animal, by hand-tracing from 40x images using the Image J freeform tool to outline the cell body. Using the analyze, ‘calculate area’ function, the program then automatically calculated the area of the hand-drawn soma in each ROI and expressed it as µm^2^.

##### Morphological Analyses

To determine measures of microglia complexity at different ages, and under different conditions, Sholl and related analyses were conducted on 20 randomly selected skeletonized microglia in the central PL of each animal. Using the overlaid contour on IBA1 sections, a sampling grid was using to help select a central ROI on each section (4-5 sections through the rostrocaudal PL/animal) (**Fig. 2B**). A 40x image of the microglia within the grid box was then captured and the microglia contained inside the grid box were assigned a number (**Fig.2C**). A random number generator (www.Calculator.net) was used to indiscriminately select 5 microglia in the grid box for tracing. A total of 20 randomly selected IBA1-positive microglia per animal were thus drawn using the 2-dimensional (2D) neuron reconstruction module (Neurolucida 360, Microbrightfield Biosciences, Williston, VT). We marked soma, process length (total dendrite length), branch points, and endpoints for each cell. A 2D convex hull for each cell was created from a line connecting endpoints (**Fig. 2D**).

Sholl analysis parameters (Neurolucida Explorer, Microbrightfield Biosciences, Williston, VT) were set to place concentric circles beginning at the center of the soma, in 5-μm intervals (**Fig.2D**). The number of intersections at each interval from the soma was used to generate a Sholl profile curve, from which several indices of microglial morphology can be derived: 1. convex hull area and, 2. number of branching nodes, 3. number of endpoints, and 4. mean process length. The 2D convex hull area is the area enclosed around the planar shape of the tracing, and convex hull perimeter is the distance around the most distal points (endpoints) that form the convex hull. The total number of nodes and total number of endpoints in the shell were also calculated. The total length in µm of all processes passing through a shell was calculated as the mean process length. All these data were exported into an Excel file and averages were calculated first for each animal, and then for the group.

### Statistics

Statistical tests were run with Prism 10 statistical analysis software (GraphPad). Data was statistically analyzed between the maternally reared infant versus adolescent animals using an unpaired t-test. Comparison of the maternally reared animals versus 1-week and 1-month separated animals within infant and adolescent categories was statistically analyzed using an ordinary one-way ANOVA comparing each experimental condition to the MR control, with Bonferroni correction. Lastly, differences amongst both experimental condition and age were accessed using a two-way ANOVA with Tukey’s multiple comparisons tests. Pairwise comparisons were considered significant if the p-value associated with them was < 0.05. Interactions between infant-adolescent relationships and experimental condition (MR, 1-WS, 1-MS) were also run using two-way ANOVA to understand the influence of experimental condition on infant-adolescent group comparisons.

## Results

### Microglia shift morphology in PL between infancy and adolescence

To provide an overview of microglia population characteristics, we first characterized how the global distribution of PL microglia differs between maternally reared infant and adolescent macaques (**Fig. 3A, B**). The density of PL microglia was significantly increased in the adolescents (mean= 394,923 +/- 19,771/ µm^2^ *10^6^) compared to infants (mean= 250,463 +/- 14,324/ µm^2^ *10^6^, F (3,3) = 1.905, (*p* <0.0001) (**Fig.3C**). However, the spacing index, which measures the clustering of neighboring microglia, was similar between maternally reared infants (0.5039 +/- 0.02052 AU) and adolescents (0.5003 +/- 0.01494 AU), F (3,3) = 1.886, (*p*=0.3945) (**Fig. 3D**). This indicated that, despite the increase in microglia density with development, when corrected for density, microglia are evenly distributed in maternally reared infant and adolescent PL.

**Figure 3.**
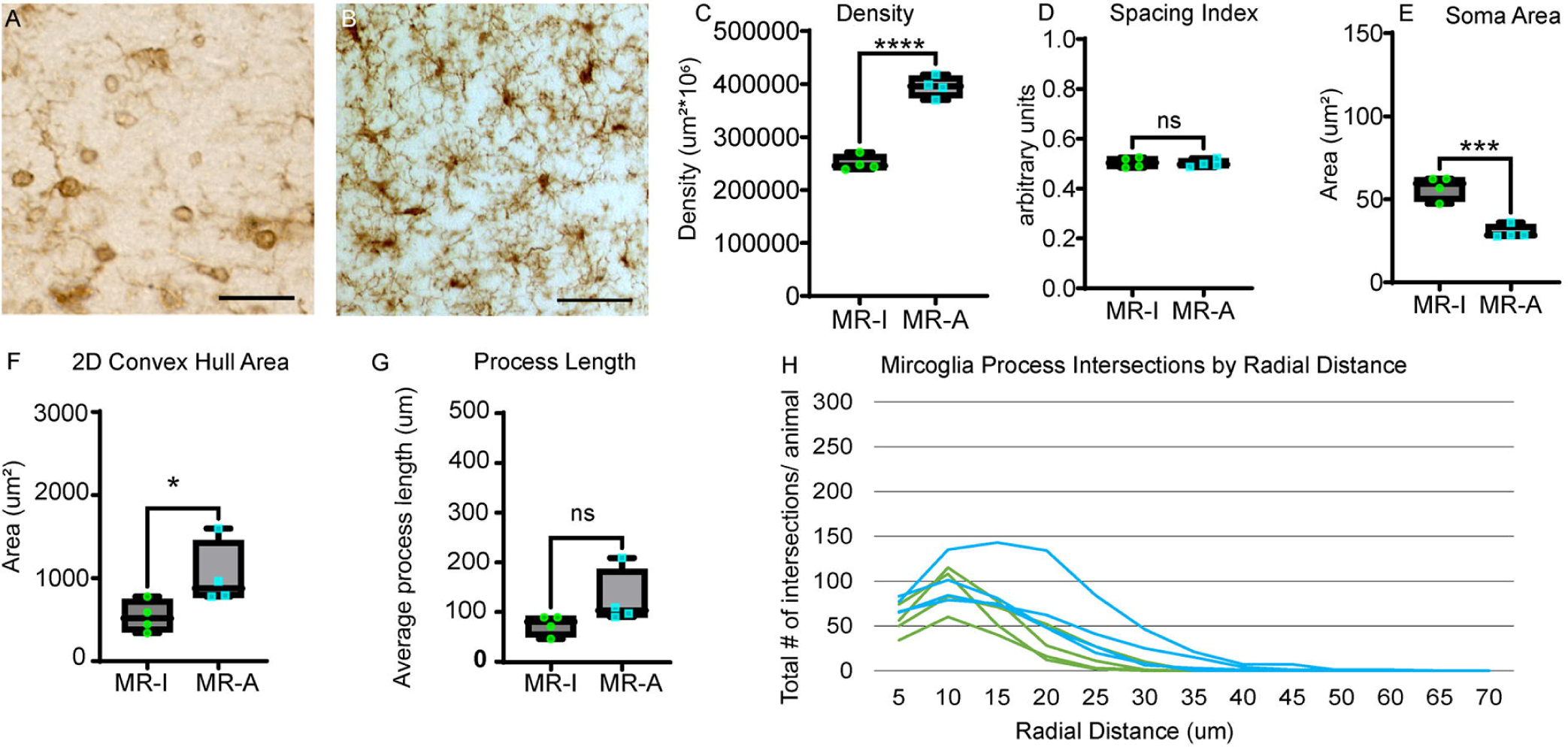
Morphological characteristics of microglia: maternally reared infants and adolescents. A. Representative IBA1-IR microglia (Infant PL). B. Representative IBA1-IR microglia (Adolescent PL). Bars = 25µm. C-G. Green= maternally reared infant; Blue= maternally reared adolescent. C. Average microglia density. D. Average Spacing Index E. Average Soma Size F. 2D Convex Hull Area. G. Average mean process length H. Microglia process intersections by radial distance for each animal.

We then focused on the morphological features of the microglia. The soma area of microglia was significantly greater in maternally reared infants (mean = 57.14 +/- 7.088 µm^2^) compared to maternally reared adolescents (mean=30.15 +/- 3.944 µm^2^, F (3,3) = 3.230, *p*=0.0003) (**Fig. 3E**), consistent with qualitative observations as seen in **Fig. 3A, B**. We also found that convex hull area is significantly greater in the adolescents (1032 +/- 385.4 µm^2^) compared to the infants (536.7 +/- 190.3 µm^2^), F (3,3 =4.104 (*p*=0.0304) (**Fig. 3F**), as is the convex hull perimeter (infants = mean 92.38+/- 14.23, adolescents = mean 130.6 +/- 19.46) F (3,3 = 1.871 (*p*=0.0097). The mean total process length, which measures all the processes passing through the convex hull, showed a trend to being greater in the adolescents (127.5 +/- 55.39) in comparison to the infants (75.08+/- 20.39) F (3,3) = 7.379 (*p*=0.0630), but did not reach significance (**Fig.3G**)

To examine the arbor complexity of microglia in individual animals in each group, we mapped the frequency of intersections between the processes and spheres at 5µm incremental radii from the soma (**Fig. 3H**). Maximal intersections in infant (green) and adolescent (blue) maternally reared animals were greatest at 10 μm from the soma. In addition, the mean number of intersections were not different between the infants (12.46 +/- 3.347) and adolescents (20.46 +/- 7.846), F (3,3) =5.521, p= 0.0552). Consistent with convex hull findings, microglia arbors ended ∼20-25 µm from the soma in infant PL, whereas in the adolescent PL, intersections were further extended out to ∼35-40 µm radial distance, supporting an overall greater distal elongation of processes in the adolescent PL. There were no differences in the mean number of endpoints, (infants= mean 6.888+/- 2.171, adolescents = mean 7.575+/- 3.084) F (3,3) = 2.018 (*p*= 0.3640) or nodes (infants = mean 2.575 +/- 1.487, adolescents = mean 2.838 +/- 1.554), (F3,3) =1.092, *p*=0.4076) in the maternally reared animals.

### Maternally separation alters microglia morphology in infants and adolescents

We then compared changes in microglia from maternally reared (MR) animals with those from the 1-WS (animals separated beginning at 1 week of life) and 1-MS (animals separated beginning at 1 month of life) animals for each age cohort.

#### *Infants* (**Fig. 4A-G**)

Overall, there were no differences in density between the maternally reared infants (264,265 +/- 14,799 /µm^2^ *10^6^) and the 1-WS infants (204,265 +/- 57,934 /µm^2^ *10^6^ *p*=0.6368) or the 1-MS infants (270,904 +/- 88,981/ µm^2^ *10^6^ *p*>0.9999) F [2, 9] = 1.218 *p* =0.3402. (**Fig. 4B**). However, the spacing index was markedly decreased in both the 1-WS (0.2146+/-0.01376 AU *p*<0.0001) and 1-MS (0.1931 +/-0.02913 AU p <0.0001) infant groups F (2, 9) = 247.7 (*p*<0.0001) in comparison to the maternally reared (MR) infant group (0.5039 +/- 0.02052 AU) (**Fig. 4C**). This suggested a more clustered distribution, despite similar densities between maternally reared and maternally separated infants.

**Figure 4.**
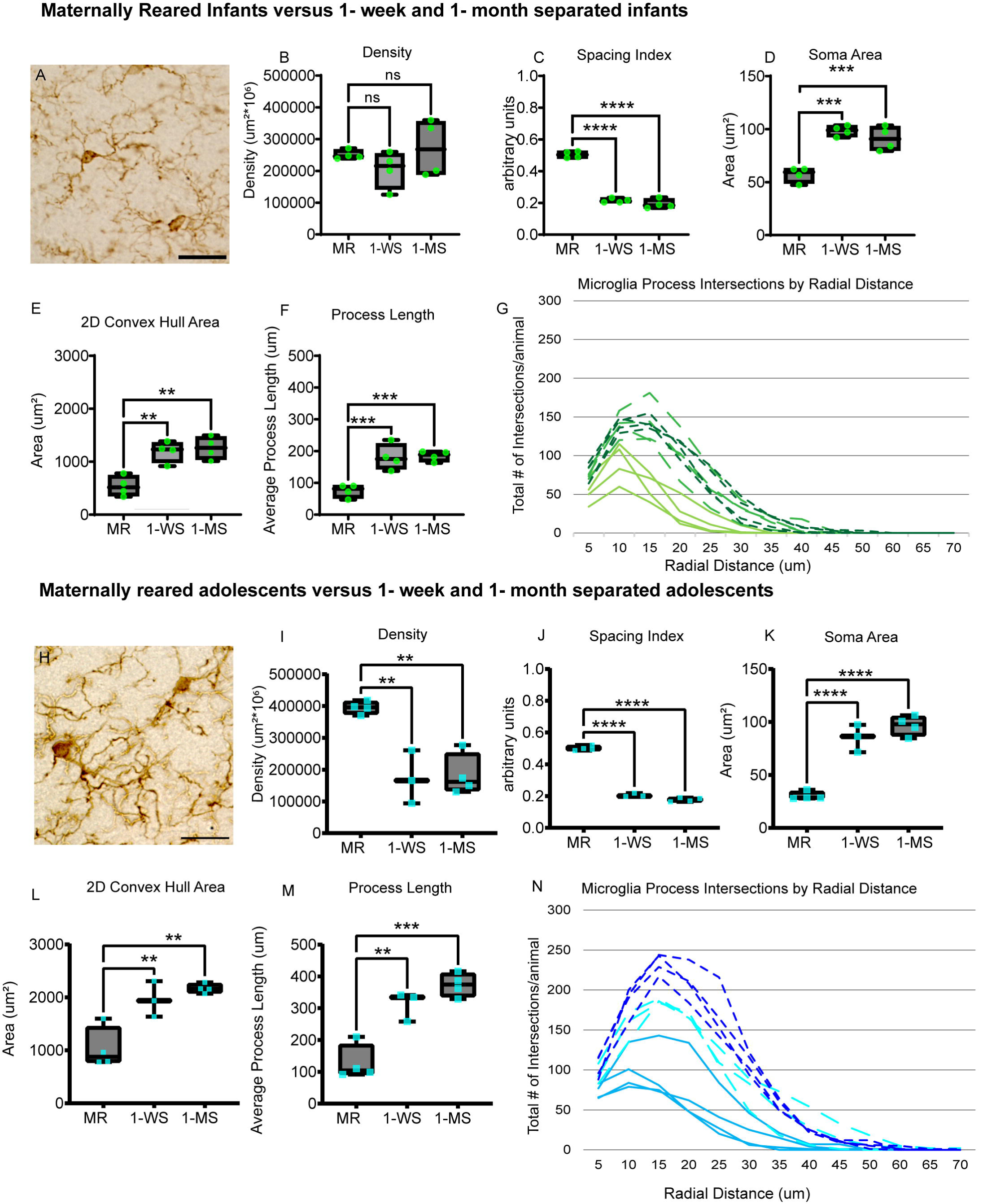
Maternal separation effects within each age cohort. Representative photomicrographs of PL microglia in A. 1-week maternally separated infants (green) and H. adolescents (blue). Scale bar = 25µm. Infant effects (B-G) and Adolescent effects (I-N). (B, I) Average microglia density; (C, J). Average Spacing Index; (D, K). Average Soma Size; (E, L) 2D Convex Hull Area; (F, M). Mean process length; (G, N). Microglia process intersections by radial distance for each animal. G. Microglia process intersections by radial distance for each animal (infants, green; adolescent, blue. MR= solid line; 1-WS= light colored dashed line; 1-MS= dark colored dashed line)

Soma area was increased in both the 1-WS (mean 98.40 +/-4.975 µm^2^ *p*=0.0001) and 1-MS (mean 91.12 +/- 11.30 µm^2^ *p*= 0.0005) infant groups compared to the MR group (mean 57.14 +/- 7.088 µm^2^) F (2, 9) = 28.70 *p*=0.0001 (**Fig. 4D**). Additionally, the convex hull area in infants was markedly increased in both 1-WS (mean= 1191+/- 196.7 µm^2^ *p=* 0.0024) and 1-MS (mean= 1257+/- 210.7µm^2^ *p*= 0.0013) groups compared to MR infants (mean= 536.7 +/- 190.3 µm^2^) F (2, 9) = 15.96 p=0.0011 (**Fig. 4E**). As expected, convex hull perimeter was similarly increased in the 1-WS (mean 136.1 +/- 10.9 um *p=*0.0022) and 1-MS (mean=138.3 +/- 13.95um *p*=0.0016) groups in comparison to the MR infants (92.38 +/- 14.23 um) F (2, 9) = 15.60 *p*= 0.0012. Consistent with a larger convex hull area and perimeter, the mean process length was increased in both 1-WS (mean= 180.7 +/- 40.55 um *p*= 0.0008) and 1-MS (mean=184.6 +/- 16.14 um *p*= p=0.0005) conditions in comparison to the MR infants (mean=75.08 +/- 20.39 um) F (2, 9) = 19.95 *p*=0.0005 (**Fig. 4F**).

The mean number of intersections in the 1-WS (28.05+/- 5.203 *p*=0.0008) and 1-MS (29.11+/- 3.271 *p*=0.0005) infant groups were more than double the mean number of intersections in the infant MR group (12.46+/- 3.347) F (2, 9) = 21.29 *p*= 0.0004 (**Fig.5H**). When the total number of intersections by distance from the soma were plotted for each animal, it was observed that microglial arbors in maternally reared infant PL extended to 25-30 µm, while in both the 1-WS (light green fragmented line) and 1-MS (forest green fragmented line) infants, arbors extended maximally between 35-45 µm away from the soma (**Fig. 4G**).

**Figure 5.**
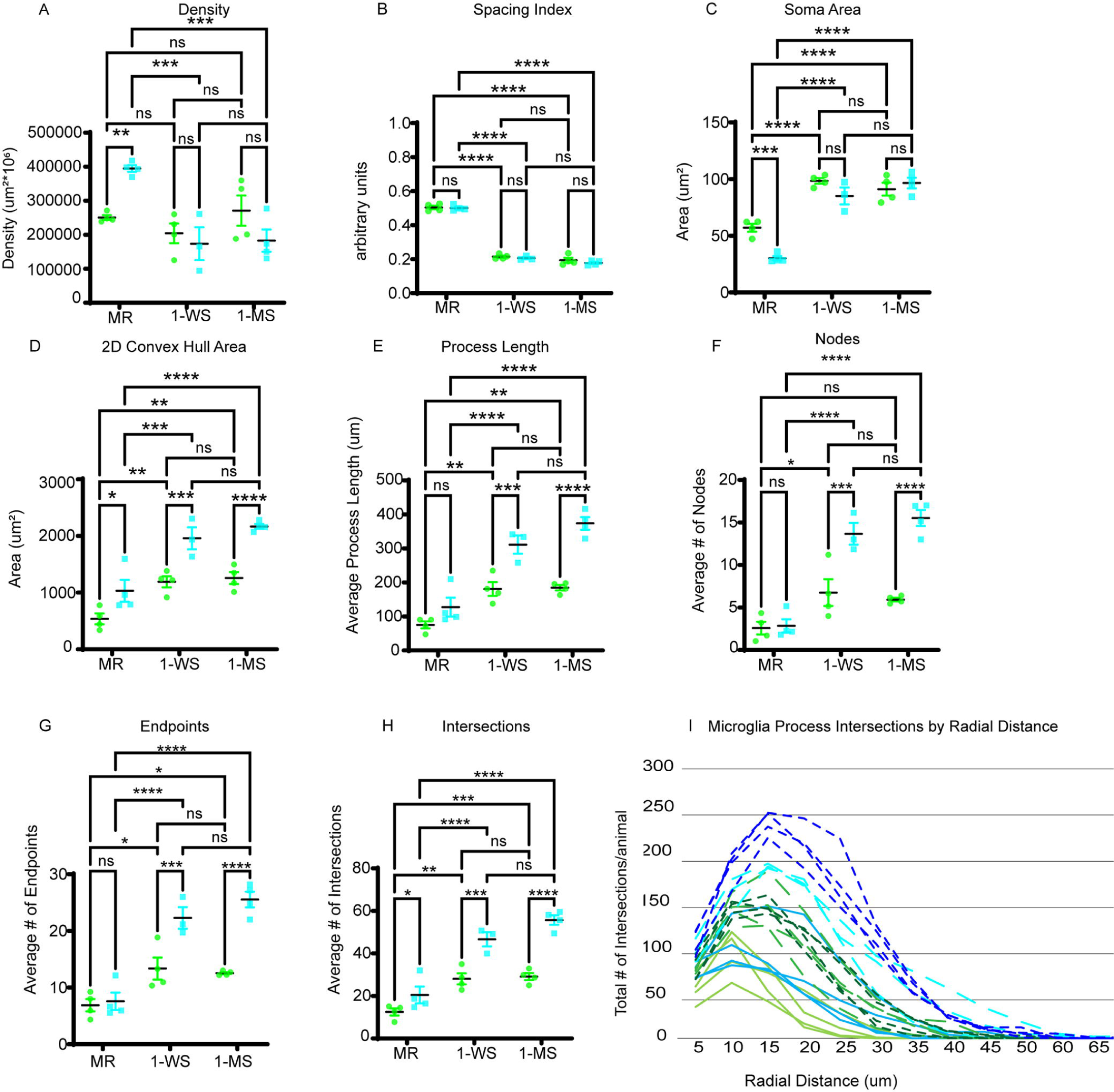
Maternal separation effects on microglial developmental trajectories. Microglia in infants (green) and adolescents (blue) in all conditions are compared using the following measures: (A) Density (B) Spacing Index (C) Soma Area. (D) 2D Convex Hull Area (E) Mean process length. (F) Average number of Nodes. (G) Average number of Endpoints. (H) Average number of Intersections. (I) Microglia process intersections by radial distance (infants, green; adolescent, blue. MR= solid line; 1-WS=dashed lighter colored line; 1-MS= dashed darker colored line).

Maternal separation also resulted in significant increases in mean process length, mean number of endpoints and nodes in infant PL microglia. The mean process length increased between the MR (75.08 +/- 20.39 µm) and both the 1-WS 180.7+/- 40.55 µm (*p*= 0.0009) and 1-MS 184.6+/- 16.14 µm (*p*=0.0007) [F (2, 9) = 19.95 *p*= 0.0005] infant animals. The mean number of endpoints also increased between the MR (6.888 +/- 2.171) and both the 1-WS 13.36 +/- 3.873 (*p*= 0.0123) and 1-MS 12.54 +/- 0.3816 (*p*= 0.0252) [F (2, 9) = 7.505 *p*= 0.0121] infant animals. There was a significant increase in the mean number of nodes between the MR (2.575 +/- 1.487) and the 1-WS (6.750+/- 3.159 *p*= 0.0348), but not the 1-MS (5.925+/- 0.4291*p*= 0.0891) [F (2, 9) = 4.74 *p*=0.0392] infant animals (**Fig. 5F-H**).

#### *Adolescents* (**Fig. 4H-N**)

The overall density of PL microglia was markedly decreased in the 1-WS (mean= 17,3621 +/- 83675/ µm^2^ *10^6^ p= 0.0024) and 1-MS (mean 18,2674 +/- 65292/ µm^2^ *10^6^ p=0.0019), when compared to the MR adolescents (mean 394,923 +/- 19,771 µm^2^ *10^6^) F (2, 8) = 17.03 *p*=0.0013 (**Fig. 4I**). The spacing index was also markedly reduced in both the 1-WS (0.2058 +/- 0.01126 AU p<0.0001) and 1-MS (0.1776 +/- 0.01240 AU *p*<0.0001) adolescent groups compared with the MR adolescents (0.5003 +/- 0.01494 AU) F (2, 8) = 713.3 *p*<0.0001 (**Fig.4J**). Together, this indicates that the microglia of maternally separated adolescents were less dense and more highly clustered, even when accounting for the decrease in density.

Soma size was also enlarged in both the 1-WS (mean=85.07 +/- 12.92 µm^2^ *p* <0.0001) and 1- MS (mean= 96.59 +/- 9.280 µm^2^ *p* <0.0001) groups compared to the MR adolescents (mean= 30.15+/- 3.944 µm^2^) F (2, 8) = 61.70 *p* <0.0001 (**Fig. 4K**). Additionally, larger 2D convex hull area was also seen in both the 1-WS (mean=1958 +/- 335.5 µm^2^ *p*= 0.0067) and 1-MS (mean= 2166+/- 87.39 µm^2^ p=0.0012) groups in comparison to MR adolescents (mean=1032+/- 385.4 µm^2^) F (2, 8) = 16.47 *p*=0.0015 (**Fig. 4L**), with 2D convex hull perimeter measurements (not shown) also greater in the 1-WS (mean=172.5 +/- 13.90 µm *p*= 0.0087) and 1-MS (mean=178.6 +/- 3.326 µm *p*= 0.0025) groups relative to the MR adolescents (mean 130.6 +/- 19.46 µm) F (2, 8) = 13.66 *p*= 0.0026. Mean total process length was higher in the 1-WS (mean=311.1+/- 46.09 µm *p*= 0.0018) and 1-MS (mean=373.6 +/- 37.06 µm *p*=0.0001) groups compared to the MR adolescents (mean=127.5 +/- 55.39 µm) F (2, 8) = 29.39 *p*=0.0002 (**Fig. 4M**).

With respect to mean total intersections in the Sholl, the 1-WS (mean= 55.39+/- 5.811 *p*=0.0012) and 1-MS (mean= 55.68 +/- 4.445 *p*< 0.0001) adolescent groups have more than double the mean intersections found in the MR adolescents (mean= 20.46 +/- 7.864, F (2, 8) = 33.82 *p*=0.001). In the frequency curves, microglia in the 1-WS and 1-MS adolescent animals had more intersections at all distances compared to MR (cyan fragmented lines compared to royal blue fragmented lines) (**Fig. 4N**). The most distal intersections of the microglia processes were slightly lower in the maternally reared (∼40 µm), compared to 1-WS (∼50 µm), and 1-MS (∼55 µm) adolescent animals.

Maternal separation also resulted in significant increases the mean process length, mean number of endpoints and nodes. The mean process length increased between the MR (127.5 +/- 55.39 µm) and both the 1-WS (311.1 +/- 46.09 µm p=0.0018) and 1-MS (373.6 +/- 37.06 µm, *p*= 0.0001) [F (2, 8) = 29.39 *p*= 0.0002] adolescent animals. The mean number of endpoints more than tripled between the MR (7.575 +/- 3.084) and both the 1-WS (22.28 +/- 3.299 *p*= 0.0004) and 1-MS (5.54 +/- 2.754 *p* <0.0001) [F (2, 8) = 39.26 *p*<0.0001] adolescent animals.

The mean number of nodes dramatically increased between the MR (2.838 +/- 1.554) and both the 1-WS (13.67+/- 2.226 *p*= 0.0001) and 1-MS (15.53+/- 1.891 *p*<0.0001) [F (2, 8) = 52.50 *p*<0.0001] adolescent animals (**Fig. 5F-H**).

Lastly, there were no statistical differences in any measures between the 1-WS and 1-MS animals in either infant or adolescent cohort: Infants: density (*p*=0.2913), spacing index (*p*=0.2534), soma size, (*p*= 0.4682), 2D convex hull area (*p*= 0.8391), mean process length (*p*= 0.9894), intersections (*p*= 0.9633), nodes (*p*= 0.8039) and endpoints (*p*=0.9088) and Adolescents: density (*p*= 0.9791), spacing index (*p*= 0.1423), soma size (*p*= 0.2124), 2D convex hull area (*p*= 0.6617), mean process length (*p*= 0.5130), intersections (*p*= 0.3821), nodes (*p*= 0.2539), and endpoints (*p*=0.3043).

### Maternal separation disrupts the normal developmental trajectory of microglia

To understand developmental effects of maternal separation on PL microglia, we analyzed data for all measures across all infant and adolescent groups (**Fig.5A-I**). Maternal separation resulted in altered normative trajectories in microglial organization and morphology. The normal increase in microglia density with adolescence was blunted by maternal separation (age x experimental condition, [F (2, 17) = 7.940 *p*=0.0037]) (**Fig. 5A**). The microglial spacing index remained similar between infancy and adolescence in all groups but was uniformly decreased in both infant and adolescent separation groups, [F (2, 17) = 17.79 *p*<0.0001]), indicating that maternal separation is associated with increased clustering that persists into adolescence (**Fig. 5B**). The pattern of normal reduction in soma size between infancy and adolescence was also lost in 1-WS and 1-MS animals, accounted for by overall increases in soma size in all separated animals compared to their MR controls, [F (2, 17) = 7.234 *p*=0.0053]) (**Fig. 5C**).

Both age and maternal separation influenced the morphological complexity of microglia (**Fig. 5D-I**). The 2D convex hull, which normally increases with age, maintained this shift. Although maternal separation increased the 2D convex hull area and perimeter further at both ages in both separated groups, development was ultimately the driving factor contributing to the differences observed in arbor size indicated by the statistically not significant interaction, [F (2, 17) = 1.429 *p*= 0.2670]; perimeter [F (2, 17) = 0.03951 *p*= 0.9613) (**Fig. 5D**).

For the mean process length (**Fig. 5E**), nodes (**Fig. 5F**), and endpoints (**Fig.5G**), there were no apparent changes with development in the MR control animals. However, maternal separation coupled with development resulted in significant increases in these three measures in both infant and adolescent groups (mean process length [F (2, 17) = 6.504 *p*= 0.0080], nodes [F (2, 17) = 12.02 *p*= 0.0006], endpoints [F (2, 17) = 10.05 *p*= 0.0013]). The increases in these measures were also exacerbated in adolescent animals compared to infants (mean process length [F (1, 17) = 60.51 *p*<0.0001], nodes [F (1, 17) = 46.51 *p*=<0.0001], endpoints [F (1, 17) = 41.38 *p*<0.0001]), indicating that maternal separation intensifies microglial complexity during development.

The expansion in microglial complexity with development was also reflected in increases in the mean number of intersections between infancy and adolescence in maternally reared and maternally separated groups [F (2, 17) = 6.416 p= 0.0084] with maternal separation resulting in additional increases in process intersections in both age groups [F (1, 17) = 66.08 *p*<0.0001] (**Fig.5H**). This was also apparent in the frequency graphs of intersections by radial distance. In general, adolescent groups (blue) have larger arbors with intersections further extended away from the soma in comparison to the infant groups (green) (**Fig.5I**) with additional increases due to maternal separation in both the 1-WS (dashed light blue) and 1-MS (dashed dark blue) adolescents. Together, patterns for convex hull, mean process length, nodes, endpoints, and intersections all suggest that the natural increase in microglia complexity with age is aberrantly heightened by maternal separation.

## Discussion

The PL is a newly recognized source of cellular plasticity in the amygdala. We hypothesized that the morphology of PL microglia embedded among immature neurons would track shifts in neuronal growth during development. We also hypothesized that maternal separation is a potent environmental event that could dynamically alter this trajectory, with implications for neural growth and circuit formation.

### Normal PL development

Immature glutamatergic neurons of the PL are likely to depend on microglia for synapse formation, like developing neurons in other brain regions (Streit, 1993; Paolicelli et al., 2011). We investigated multiple measures of microglia morphology in the PL of both infant and adolescent animals to gain insight into functional state. In maternally reared animals, the PL microglia population density increased between infant and adolescent animals and maintained a homogenous distribution. At the same time, microglia soma size decreased, while their overall arbor area increased between infancy and adolescence. These findings are consistent with the typical developmental transition of microglia from an “immature” to “mature” phenotype (Savage et al., 2019; Dos Santos et al., 2020; Vidal-Itriago et al., 2022).

Prior literature has characterized the predominant microglial morphology during infancy having enlarged soma size and minimal processes (‘amoeboid’) (Ivy and Killackey, 1978; Innocenti et al., 1983). In this phenotype, microglia scavenge and phagocytose whole cells and have a high migratory capacity (see review (Vidal-Itriago et al., 2022)). By adolescence, the distinctly smaller soma size, and elongated processes, appear closer to a ‘ramified’ phenotype (Vidal-Itriago et al., 2022). This transition is consistent with the morphological shifts we observed in the microglia of maternally reared animals in the PL. Between infancy and adolescence, there was a substantial decrease in microglial soma size, along with an increase arbor size. While microglia in the ramified phenotype do not typically engage in whole cell phagocytosis, they are thought to surveil their surrounding parenchyma, and phagocytose synaptic elements, a process known as synaptic pruning (Tremblay et al., 2010). During brain development, microglia interface with activity-dependent changes in synapse formation to eliminate weakened synapses (Whitelaw et al., 2023). Since the PL is packed with immature glutamatergic neurons that gradually mature between infancy and adolescence (presumably to integrate into amygdala circuitry) (McHale-Matthews et al., 2023), it is notable that microglia density increases, and morphology shifts to a more ramifying (pruning) phenotype, during this time.

### Maternal separation effects on PL

Maternal separation is a highly potent environmental disruption for young animals, including macaques and humans (Seay et al., 1962; Seay and Harlow, 1965; Coe et al., 1988; Fabricius et al., 2008; Koe et al., 2016; Cameron et al., 2017). Surprisingly, maternal separation had a large effect not only on the PL microglia in infants (separated from the mothers for a few months), but also on PL microglia in adolescents who had been separated from the mother many years prior. The effects of maternal separation were much larger than the typical age-related morphological changes observed in microglia between infancy and adolescence (**Fig. 6**). Microglia in the maternally separated animals at both ages had relatively enlarged somas and increased number of processes and mean process length, characteristics consistent with a ‘hyper-ramified’ phenotype (Hinwood et al., 2013; Smith et al., 2019). The maintenance of this striking phenotype over years in the separated adolescents suggests that the effects of early maternal separation are long-lasting. Thus, early environmental influences, such as stress, that stimulate synaptic activity may alter developing neural circuitry in concert with microglia activity (Catale et al., 2020; Block et al., 2022).

**Figure 6.**
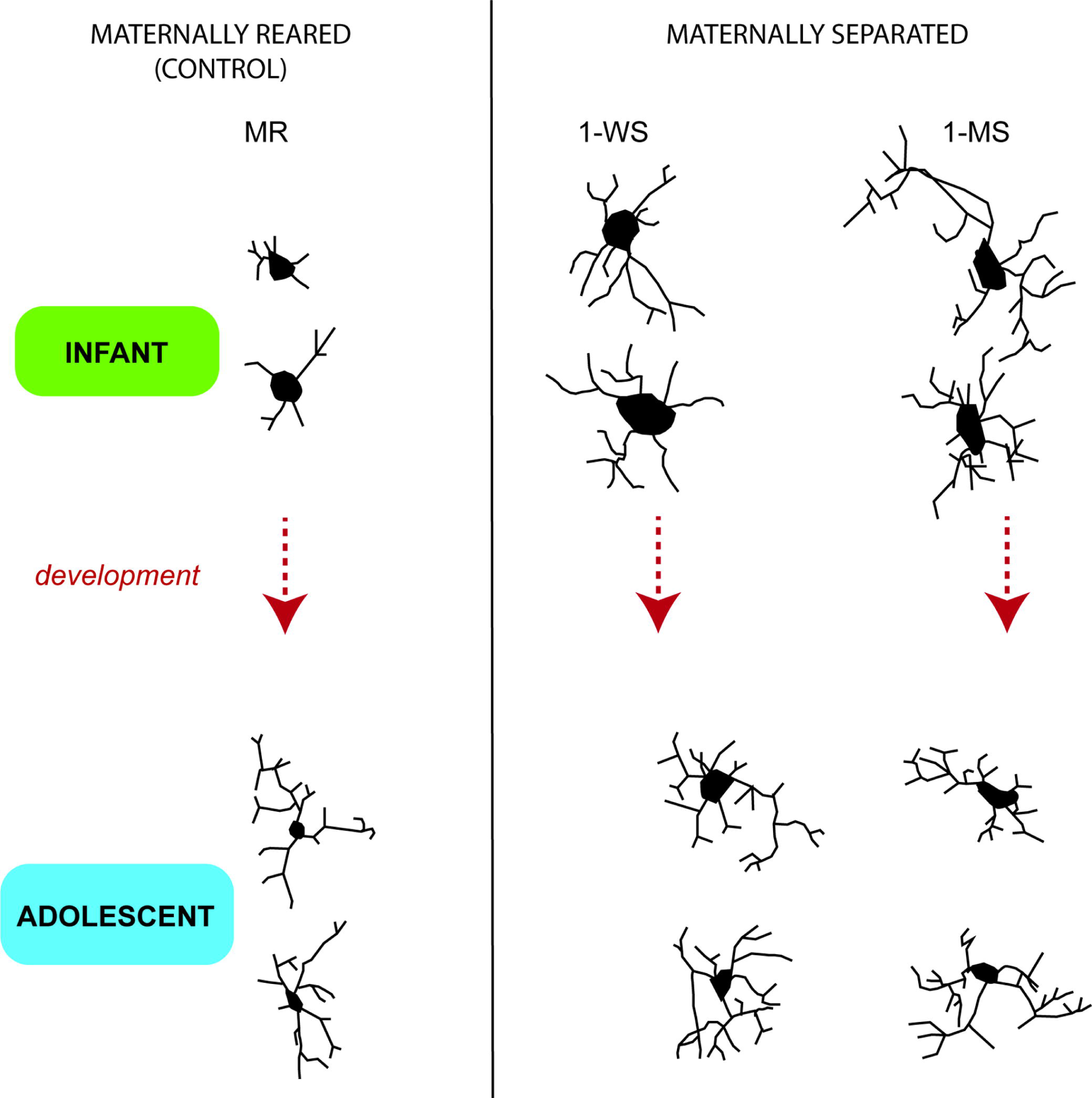
Camera lucida drawings of typical microglia in maternally reared infants and adolescents compared to 1-WS and 1-MS in each cohort. Drawn with 40x objective.

### A persistent hyper-ramified microglia phenotype

The ‘hyper-ramified’ phenotype, distinguished by increased microglial soma area, process length and complex branching patterns (increased nodes, intersections, and endpoints) is observed when animals are subjected to chronic stress (Hinwood et al., 2012; Hinwood et al., 2013; Smith et al., 2019). The hyper-ramified subtype has been conceptualized as an ‘intermediate phenotype’ (Streit, 1996; Walker et al., 2014), that arises before the emergence of the ‘hypertrophic phenotype (a ‘bushy’ morphology with retracted, thickened dendritic processes and large soma) observed in classic inflammation (Streit, 1996). More recent work indicates that the effects of stress on microglial ramification can also occur without accompanying evidence of inflammation (e.g. increases in interleukin-1beta, MHC-II, CD68, terminal deoxynucleotidyl transferase dUTP nick end labeling, and activated caspase-3) (Smith et al., 2019). Interestingly, rodents bred to be ‘low responders’ to novelty (novelty-averse) have more ‘hyper-ramified’ microglia than ‘high responders’, suggesting that in the non-challenged state, animals that are inherently stress-sensitive have altered microglial phenotypes (Maras et al., 2022). In humans, the ‘hyper-ramified phenotype’ is among the diverse microglial phenotypes commonly found in the normal adult brain (as assayed in the anterior cingulate). However, in the brains of people who suffered from depression and died by suicide, the prevalence of ‘hyper-ramified’ microglia is significantly increased (Torres-Platas et al., 2014b; Torres-Platas et al., 2014a).

### Microglia-neuron interactions following stress

Microglial shifts in the face of stress have recently become viewed as adaptive homeostatic responses, which correlate with the duration and severity of the stressor (Walker et al., 2014; Woodburn et al., 2021). These responses are termed ‘para-inflammation’ since classic neuroinflammatory responses such as macrophage activation, pro-inflammatory cytokine production, leukocyte recruitment, and tissue damage may not be present. During para-inflammation, the immune system sustains low activation to support homeostatic function. A proposed mechanism for ‘para-inflammatory responses’ is the increased activity in glutamatergic neurons during stress and the increase in glucocorticoids, which in turn triggers compensatory responses in microglia along with shifts in their morphological phenotype (Woodburn et al., 2021). Accordingly, the hyper-ramified phenotype is functionally associated with experience-dependent modifications in several brain regions in rodent models, and can impact synaptic formation and maintenance (Hinwood et al., 2013; Wohleb et al., 2018; Smith et al., 2019; Bollinger et al., 2022). In this model, increased glutamate release induced by chronic stress facilitates microglial synaptic phagocytosis, which can be seen as a protective (adaptive) mechanism. Here, our finding of a shift to a more hyper-ramified microglia phenotype in the PL of maternally separated animals in both infants and adolescents could reflect protective microglial responses to facilitate neuronal survival, in this densely populated region of immature glutamatergic neurons.

Supporting the idea compensatory neuronal-microglial interactions in responses to stress, we recently showed in this cohort of infant monkeys PL-specific decreases in neuronal genes associated with spinogenesis, axogenesis, and migration during maternal separation (de Campo et al., 2017). For example, mRNA for the activity-dependent gene TBR1 which is required for glutamatergic spine formation (Darbandi et al., 2018), was strongly downregulated in the maternal separation condition. As TBR1 is normally highly expressed in the PL, its decrease at the mRNA level in maternally separated infants may reflect synapse loss due to microglial hyperramification/phagocytosis, and loss of excitatory transmission. Alternately, reduction of TBR1 may also be a cell autonomous protective mechanism. More exploration of gene expression changes in both infant and adolescent PL is clearly needed to understand PL neuron-microglial interactions after maternal separation.

One of the most surprising findings of our study was that ‘hyper-ramified’ microglia were a dominant phenotype in maternally separated adolescent animals, many years after their separation. Although the mechanism of this persistent phenotype is unclear, the phenomenon is reminiscent of an exacerbated ‘hyper-ramified’ phenotype in individuals with history of major depression (Torres-Platas et al., 2014b; Torres-Platas et al., 2014a).The idea that ‘priming’ by pre-existing stress can have long-lasting effects on microglia function and behavior has been demonstrated in rodent models and is important for understanding later-developing psychiatric and neurologic illnesses (Hammond et al., 2018; Catale et al., 2020).

### Limitations

There are a few inherent limitations to this study, most driven by the value and labor involved in studying the macaque species such as small sample sizes due to the sheer value of the primate animals used and their prolonged gestational period. Although we were able to determine strong changes in microglial morphology, a larger sample may have permitted detection of subtle differences between the 1-WS and 1-MS conditions. A total of 19 female and 4 male monkeys were utilized in this study. In future work, sex differences in the effects of maternal separation on PL microglia in male and female macaques should be explored. Regrouping of all adolescent animals in the last year of life may have increased stress, with unknown effects on microglial morphology. Nonetheless, clear shifts in microglia in the infant cohort by maternal separation were similar, and frequently enhanced, in the adolescent separation group, and there was little effect on adolescent maternally reared animals. This suggests that while later ‘re- grouping’ stress (as well as other intervening life experiences) may have affected microglia in adolescent PL, maternal separation was required for the ‘hyperramified’ phenotypes observed. Finally, conclusions on what is functionally occurring with the microglia cannot be solely based on their morphology. Although we have identified dynamic developmental morphologically differences between our experimental groups, future studies will be needed to investigate the phagocytic state of the microglia and the engulfment of putative synapses.

## Conclusion

In conclusion, our results show that maternal separation induced an altered developmental trajectory of the microglia in the macaque PL. These changes persist in adolescence along with ongoing behavioral abnormalities. Because environmental challenges and activity influences have been shown to interfere with the development of microglia including their phagocytic activity, microglial changes due to maternal separation may in turn also alter PL neuronal maturational processes thought to support emerging amygdala function in young animals.

## Acknowledgements

Nannette Alcock, for histological assistance. This work was funded through the John D. and Catherine T. MacArthur Foundation Network on Early Experience and Brain Development (JLC), The Schmitt Program for Integrative Neuroscience at the Del Monte Institute for Brain Science (URMC), the National Institute of Neurological Disorders (T32 NS115705 DK), the National Institute of Mental Health (R21MH127486, JF and JC; PO1MH41712, JC), and DHHS/PHS/NIH (P50 HD103536, DK).

